# Assaying uptake of endocrine disruptor compounds in zebrafish embryos and larvae

**DOI:** 10.1101/169987

**Authors:** J. Paige Souder, Daniel A. Gorelick

## Abstract

To study the effects of environmental endocrine disruptors (EEDs) on aquatic animals, embryos and larvae are typically incubated in water containing defined concentrations of EEDs. However, the amount of EED uptake into the animal is often difficult to determine. Using radiolabeled estradiol ([^3^H]E2), we previously developed a rapid, straightforward assay to measure estradiol uptake from water into zebrafish embryos and larvae. Here, we extend this approach to measure the uptake of two additional EEDs, bisphenol A (BPA) and ethinyl estradiol (EE2). As with E2, the uptake of each compound by individual larvae was low (< 6%), and increased with increasing concentration, duration, and developmental stage. We found that E2 and EE2 had similar uptake under equivalent exposure conditions, while BPA had comparatively lower uptake. One application of this assay is to test factors that influence EED uptake or efflux. It has been suggested that persistent organic pollutants (POPs) inhibit ABC transporters that may normally efflux EEDs and their metabolites, inducing toxicity in aquatic organisms. We measured [^3^H]E2 levels in zebrafish in the presence or absence of the POP PDBE-100, and cyclosporine A, a known inhibitor of ABC transporters. Neither chemical significantly affected [^3^H]E2 levels in zebrafish, suggesting that zebrafish maintain estradiol efflux in the presence of PDBE-100, independently of cyclosporine A-responsive transporters. These uptake results will be a valuable reference for EED exposure studies in developing zebrafish, and provide a rapid assay to screen for chemicals that influence estrogen-like EED levels *in vivo.*

## INTRODUCTION

Environmental endocrine disruptors (EEDs) are small molecules that mimic endogenous hormones. EEDs can negatively impact the health of humans and wildlife by disrupting endogenous hormone signaling (Diamanti-Kandarakis et al., 2009). Estrogen-like EEDs are a broad class of EEDs including endogenous compounds like 17-β-estradiol (E2) and synthetic compounds like bisphenol A (BPA), commonly found in manufactured plastics.

Aquatic animal models, such as zebrafish, are used to study the environmental impact of EEDs. One common approach is to expose zebrafish embryos to known and suspected EEDs and assay their toxicity (Bouwmeester et al., 2016; Carroll et al., 2014; Gorelick et al., 2014; Padilla et al., 2012; Tal et al., 2016). However, for the majority of EEDs, information regarding the precise uptake and excretion is lacking. We previously developed an assay to measure [^3^H]E2 uptake in zebrafish embryos, and found that supraphysiologic concentrations of E2 in fish water are required to achieve physiologically-relevant doses in embryos (Souder and Gorelick, 2017). We also found that E2 uptake is dependent on exposure concentration, duration and developmental stage (Souder and Gorelick, 2017). We sought to determine if this is also true for other estrogen-like EEDs, by testing the uptake of the pharmaceutical estrogen analog, ethinyl estradiol (EE2) and the non-steroidal synthetic estrogen, bisphenol A (BPA).

In addition to quantifying EED uptake, it would also be useful to identify chemicals that influence EED uptake or efflux to discover novel mechanisms of toxicity and to potentially inhibit the uptake of toxic EEDs. Major drug transporters like P-glycoprotein (P-gp) are known to regulate uptake and efflux of an array of structurally diverse substrates, including xenobiotics and pharmaceuticals (Aller et al., 2009; Ambudkar et al., 1999). Though previous efforts have investigated P-gp transport of steroids and xenobiotics *in vitro* (Kim and Benet, 2004), the degree to which P-gp influences EED efflux *in vivo* is less well understood.

Using our assay for measuring radiolabeled estradiol uptake, we quantified the uptake of [^3^H]EE2 and [^3^H]BPA in zebrafish embryos at multiple concentrations, exposure durations, and developmental stages. We found that less than 5% of EE2 and BPA are taken up following 24 hour exposure, and that EE2 and BPA uptake are dependent on concentration, duration, and developmental stage. When comparing E2 uptake to EE2 and BPA, we found that EE2 uptake is similar to E2, whereas BPA uptake is substantially lower. Additionally, we found that inhibition of *abcb4*, a zebrafish P-gp orthologue (Fischer et al., 2013), did not affect E2 uptake. Our results support the hypotheses that supraphysiologic concentrations of EEDs in water are required to achieve physiologic concentrations *in vivo.* Our results also suggest that environmentally-relevant concentrations of E2 are not influenced by the drug transporter *abcb4.*

## METHODS

### Zebrafish

Adult zebrafish were raised at 28.5°C on a 14-h light, 10-h dark cycle in the UAB Zebrafish Research Facility in a recirculating water system (Aquaneering, Inc., San Diego, CA). All zebrafish used for experiments were wildtype, AB strain (Westerfield, 2000). All procedures were approved by the UAB Institutional Animal Care and Use Committee.

### Embryo Collection

Adult zebrafish were allowed to spawn naturally in groups. Embryos were collected in intervals of 10 minutes to ensure precise developmental timing, placed in 100mm × 15mm Petri dishes at a density of no more than 100 per dish in E3B media (60X E3B: 17.2g NaCl, 0.76g KCl, 2.9g CaCl_2_-2H_2_O, 2.39g MgSO_4_ dissolved in 1L Milli-Q water; diluted to 1X in 9L Milli-Q water plus 100 μL 0.02% methylene blue), and then stored in an incubator at 28.5°C on a 14-h light, 10-h dark cycle until treatment.

### Embryo treatments

For uptake experiments, embryos were treated in tritiated ethinyl estradiol ([6,7-^3^H(N)]- 17α-ethynylestradiol, 1 mCi/mL, 60 Ci/mmol, American Radiolabeled Chemicals Inc. #ART1321, Lot #170203), tritiated bisphenol A ([ring-3H]-bisphenol A, 1mCi/mL, 25 Ci/mmol, American Radiolabeled Chemicals Inc. #ART1676, Lot #170320), tritiated estradiol ([6,7-^3^H(N)]-17β-estradiol, 1 mCi/mL, Perkin Elmer NET013250UC), or vehicle (0.1% ethanol (EtOH)) and diluted to final concentration in E3B at the time of treatment. ABC transporter inhibitor experiments were conducted using cyclosporin A (Enzo Life Sciences, #380-002-M100) or 2,2’,4,4’,6-pentabromodiphenyl ether (PDBE-100) (AccuStandard, #FF-BDE-100N, Lot #26813) dissolved in dimethylsulfoxide (DMSO), or vehicle control (0.1% DMSO), diluted to final concentration in E3B at the time of treatment. Rhodamine B (Acros Organics, ≥ 98% pure, #AC296570100) was dissolved in methanol (MeOH) and diluted to final concentration in E3B.

### [^3^H] Uptake assay

Isotopic uptake assays were performed as described previously (Souder and Gorelick, 2017). Briefly, embryos were exposed in 24-well plates to 2 mL of treatment solution per well, ten embryos per well. All embryos were manually dechorionated prior to exposure and incubated at 28.5°C on 14-h light, 10-h dark cycle unless noted. Radioactivity of individual homogenized embryos was measured using liquid scintillation counting. Background radioactivity of vehicle-control groups was negligible and is therefore excluded from graphs. [^3^H] radioactivity was converted to pmol using a standard curve generated for each chemical (Fig. S1) with a limit of detection of 0.01 pmol for each chemical. For experiments requiring addition of non-tritiated compounds, drugs were added to the well at the same time as the tritiated compound and DMSO was used as a vehicle control. No toxicity was observed at the concentrations used for treatment (not shown).

### Rhodamine B uptake assay

For rhodamine B uptake experiments, embryos were exposed as for uptake experiments, except that embryos were exposed in the dark to prevent the loss of fluorescence during treatment. At the end of the exposure period, embryos were washed three times in fresh E3B and 0.01 mg/ml tricaine was added to immobilize embryos for imaging. Anesthetized embryos were embedded in 3% methyl cellulose in E3B and imaged on a Nikon AZ100 microscope with Andor Clara digital camera. Fluorescence was quantified from whole embryos using ImageJ software (Schneider et al., 2012) by tracing the outline of the entire embryo and averaging the mean gray value of 10 embryos per experiment.

### Experimental design and data analysis

Experiments were performed on 10 embryos from a single clutch per treatment group or vehicle control group. Experiments were performed at least 3 times (n ≥ 3) using embryos from different clutches. Mean pmol uptake and mean percent uptake from each group were used for comparing treatment groups between experiments. Mean integrated density was used to compare fluorescence between RhB-treated groups. A two-tailed, unpaired Student’s t-test was used when testing for statistical significance between two groups, one-way ANOVA with Tukey’s test for multiple comparisons was used when comparing uptake between > 2 groups, and one-way ANOVA with Dunnett’s test for multiple comparisons was used when comparing fold-change in uptake of [^3^H]EE2 and [^3^H]BPA versus [^3^H]E2. Statistical significance was set at p < 0.05. GraphPad Prism 7.0a software was used for all statistical analyses and for producing graphs.

## RESULTS

### EED uptake is concentration- and duration-dependent

To test the effect of increasing concentration on chemical uptake, we exposed 48 hour post fertilization (hpf) dechorionated zebrafish embryos to three different concentrations of [^3^H]EE2 (1 nM, 5 nM, 10 nM) or [^3^H]BPA (5 nM, 10 nM, 20 nM) for one hour. Consistent with previous results measuring [^3^H]E2 uptake(Souder and Gorelick, 2017), the absolute amount absorbed in pmol of both EE2 and BPA increased with increasing exposure concentration. [^3^H]EE2 absorption increased by 11-fold between 1 nM and 10 nM exposure (Fig. 1A, 1 nM treatment = 0.015 ± 0.0032 pmol (mean ± SD), 5 nM = 0.074 ± 0.0020 pmol, 10 nM = 0.17 ± 0.017 pmol). [^3^H]BPA absorption was less robust, increasing by 3.6-fold between 5 nM and 20 nM exposure (Fig. 1D, 5 nM treatment = 0.015 ± 0.00088 pmol, 10 nM = 0.34 ± 0.0051 pmol, 20 nM = 0.054 ± 0.017 pmol).

**Figure 1.**
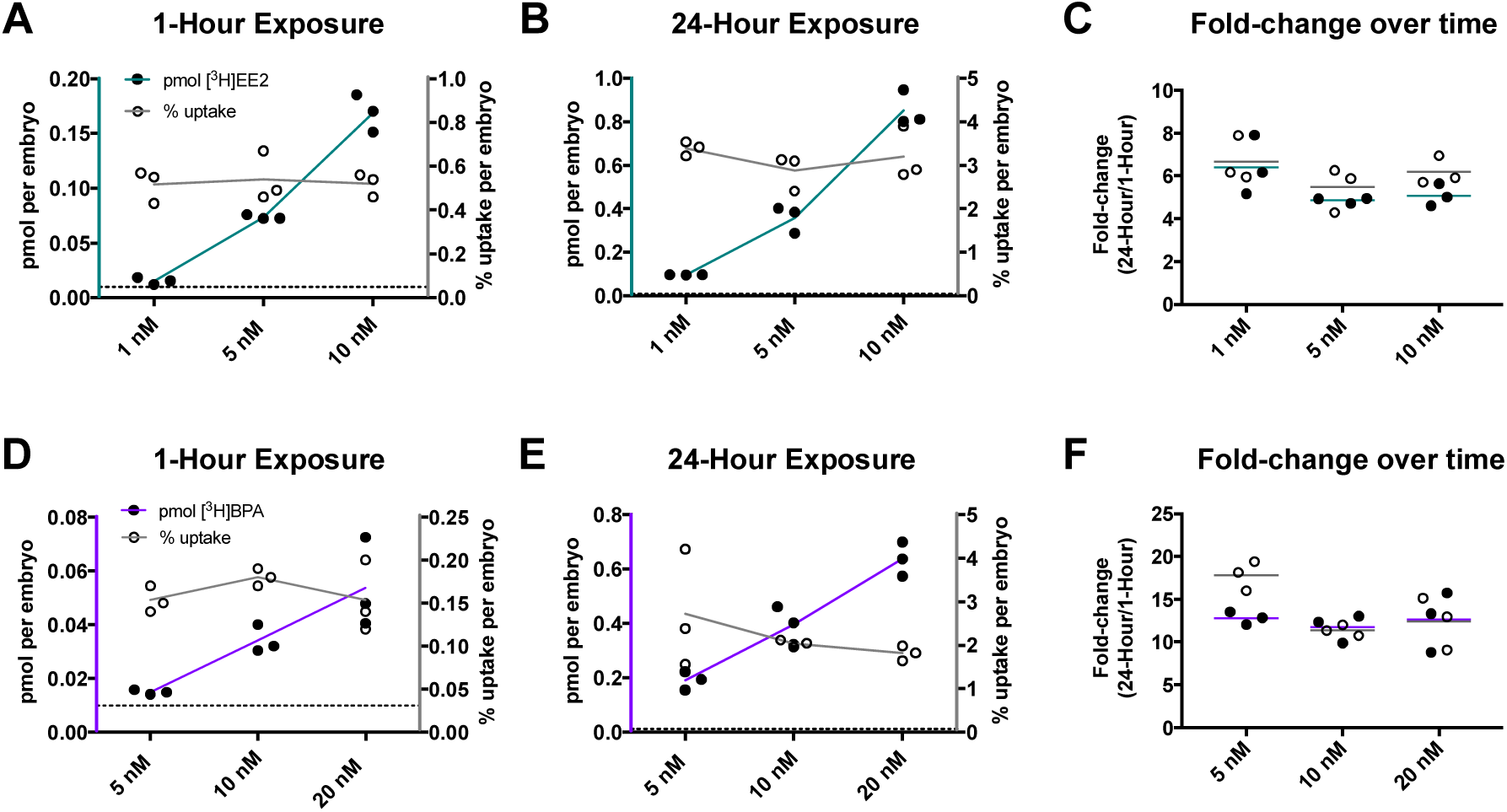
EE2 uptake is greater than BPA at multiple exposure concentrations and durations. **(A-F)** Embryos at 48 hours post fertilization were exposed to three different concentrations of [^3^H]EE2 **(A-C)** or [^3^H]BPA **(D-F)** for one hour **(A,D)** or 24 hours **(B,E)**. Radioactivity was measured using a scintillation counter and used to calculate pmol per embryo (black circles) and percent uptake per embryo (white circles). For each compound, pmol uptake increased in a dose-dependent manner (teal or purple lines connecting black circles), whereas percent [^3^H]E2 uptake remained constant (grey lines connecting white circles). Each circle represents the mean uptake from a single experiment (n=3) assaying 10 embryos per experiment. Horizontal dotted line represents the limit of detection (0.01 pmol). **(C)** Both pmol and percent [^3^H]EE2 uptake increased approximately 6-fold following 24-hour exposure compared to 1-hour exposure. Horizontal lines represent the mean fold change in pmol (teal) or percent uptake (grey). **(F)** Both pmol and percent [^3^H]BPA uptake increased approximately 13fold following 24-hour exposure compared to 1-hour exposure. Horizontal lines represent the mean fold change in pmol (purple) or percent uptake (grey).

We next tested the hypothesis that increasing exposure duration would increase EED uptake. We exposed 48 hpf dechorionated embryos to identical concentrations of [^3^H]EE2 or [^3^H]BPA for 24 hours. We found that pmol absorption increased as concentration increased. [^3^H]EE2 absorption increased by 9-fold between 1 nM and 10 nM exposure (Fig. 1B, 1 nM treatment = 0.096 ± 0.00093 pmol (mean ± SD), 5 nM = 0.36 ± 0.062 pmol, 10 nM = 0.85 ± 0.081 pmol). [^3^H]BPA absorption increased less robustly, with a 3.3-fold increase between 5 nM and 20 nM (Fig. 1E, 5 nM treatment = 0.19 ± 0.034 pmol, 10 nM = 0.40 ± 0.070 pmol, 20 nM = 0.64 ± 0.063 pmol). For each concentration, absorption increased when increasing exposure duration from 1 hour to 24 hours by an average of 5.5 ± 0.69 fold for [^3^H]EE2 (Fig. 1C), and an average of 12.4 ± 0.45 fold for [^3^H]BPA (Fig. 1F). Together, these results demonstrate that the uptake of [^3^H]EE2 and [^3^H]BPA are concentration- and duration-dependent.

We also determined the percent uptake of [^3^H]EE2 or [^3^H]BPA per embryo by dividing the embryo radioactivity by the radioactivity of the water prior to zebrafish exposure. Percent uptake remained constant at increasing concentrations for both [^3^H]EE2 (Fig. 1A-B; 1 hour exposure: 1 nM = 0.52 ± 0.076% (mean ± SD); 5 nM = 0.54 ± 0.11%, 10 nM = 0.52 ± 0.053%; 24 hour exposure: 1 nM = 3.39 ± 0.16%, 5 nM = 2.88 ± 0.41%, 10 nM = 3.12 ± 0.61%) and [^3^H]BPA (Fig. 1D-E; 1 hour exposure: 5 nM = 0.15 ± 0.015% (mean ± SD); 10 nM = 0.18 ± 0.010%, 20 nM = 0.15 ± 0.041%; 24 hour exposure: 5 nM = 2.71 ± 1.4%, 10 nM = 2.04 ± 0.080%, 20 nM = 1.81 ± 0.17%)) when holding exposure duration constant. When comparing exposure durations at each concentration, there was an average 6.1 ± 0.49 fold increase from 1 to 24 hour exposure for [^3^H]EE2 and an average 13.9 ± 2.8 fold increase for [^3^H]BPA. We conclude that percent uptake of [^3^H]EE2 and [^3^H]BPA is duration-dependent, but not concentration-dependent.

Concentrations for [^3^H]EE2 were chosen based on concentrations used for [^3^H]E2 uptake experiments. Initial experiments with [^3^H]BPA demonstrated absorption below the limit of detection (0.01 pmol) in embryos treated with 1 nM [^3^H]BPA, therefore higher concentrations (5 nM – 20 nM) were chosen for this compound. For subsequent experiments testing variables other than concentration, the 5 nM dose for [^3^H]EE2 and the 10 nM dose for [^3^H]BPA were chosen, as these concentrations were the lowest concentrations reliably above the assay limit of detection.

### EE2 and E2 uptake exceed BPA uptake at multiple concentrations and durations

We next sought to compare absorption of the estrogen-like EEDs, EE2 and BPA, to 17- β-estradiol absorption determined by our previous studies(Souder and Gorelick, 2017). When comparing pmol absorption following 5 nM exposure at 48 hpf, [^3^H]EE2 had a significantly higher absorption than [^3^H]E2 after 1 hour (one-way ANOVA, p = 0.0059), though there was no significant change in absorption with 24 hour exposure (Fig. 3A, Table S1, S2; 1 hour fold-change = 2.02 ± 0.058, 24 hour fold-change = 1.01 ± 0.18). [^3^H]BPA absorption was significantly lower at both exposure durations (Fig. 3A, Table S1, S2; 1 hour fold-change = 0.41 ± 0.025, p = 0.0014; 24 hour fold-change = 0.48 ± 0.085, p = 0.0114). Similar results were obtained when comparing 10 nM exposure at 48 hpf for [^3^H]EE2, with significantly higher absorption following 1 hour (one-way ANOVA, p = 0.0028), but not 24 hour, exposure (Fig. 3B, Table S1, S2; 1 hour fold-change = 2.26 ± 0.23, 24 hour fold-change = 1.03 ± 0.10). [^3^H]BPA uptake was also significantly lower than [^3^H]E2 at both exposure durations with 10 nM treatment (Fig. 3B, Table S1, S2; 1 hour fold-change = 0.46 ± 0.070, p = 0.0035; 24 hour fold-change = 0.48 ± 0.085, p = 0.0048).

Comparing percent uptake of 5 nM [^3^H]EE2 and [^3^H]BPA to [^3^H]E2, we did not see a significant change in percent [^3^H]EE2 uptake at either exposure duration, likely due to the increased variability in percent uptake compared to pmol uptake (Fig. 3A, Table S1, S2; 1 hour fold-change = 1.41 ± 0.30; 24 hour fold-change = 0.75 ± 0.11). We did, however, observe a significant increase following 1 hour exposure with 10 nM treatment (one-way ANOVA, p = 0.0071). There was no change in percent [^3^H]EE2 uptake compared to [^3^H]E2 uptake following 10 nM, 24 hour exposure (Fig. 3B, Table S1, S2; 1 hour fold-change = 1.77 ± 0.18; 24 hour fold-change = 0.84 ± 0.16). Similarly, we did not see a significant change in percent [^3^H]BPA uptake with 5 nM exposure, but found a significant decrease in percent [^3^H]BPA uptake with 10 nM exposure (one-way ANOVA; 1 hour exposure: p = 0.0163, 24 hour exposure: p = 0.0042)(Fig. 3A, B; Table S1, S2; 5 nM: 1 hour fold-change = 0.40 ± 0.036, 24 hour fold-change = 0.71 ± 0.35; 10 nM: 1 hour fold-change = 0.61 ± 0.035, 24 hour fold-change = 0.54 ± 0.025).

### EE2 and BPA uptake are age-dependent

Studies of EED toxicity expose embryos at multiple developmental stages, beginning at early embryonic stages and throughout organogenesis, which is largely complete by 5 dpf. To test EED uptake as a function of age, we exposed embryos beginning at five different developmental stages between 6 and 96 hours post fertilization (hpf) to a single concentration of [^3^H]EE2 (5 nM) or [^3^H]BPA (10 nM) for 1 hour. For [^3^H]EE2, mean pmol uptake when starting treatment at 96 hpf was 2.8-fold higher than when starting treatment at 6 hpf (Fig. 2A, Table S3; 6 hpf treatment = 0.037 ± 0.007 pmol (mean ± SD), 24 hpf = 0.038 ± 0.007 pmol, 48 hpf = 0.074 ± 0.002 pmol, 72 hpf = 0.072 ± 0.009 pmol, 96 hpf = 0.10 ± 0.006 pmol). There was not a statistically significant difference when comparing treatment of 6 vs. 24 hpf embryos (one-way ANOVA, p = 0.9996) or 48 vs. 72 hpf embryos (p = 0.9989), but other comparisons showed a significant increase in older versus younger embryos (Table S3).

**Figure 2.**
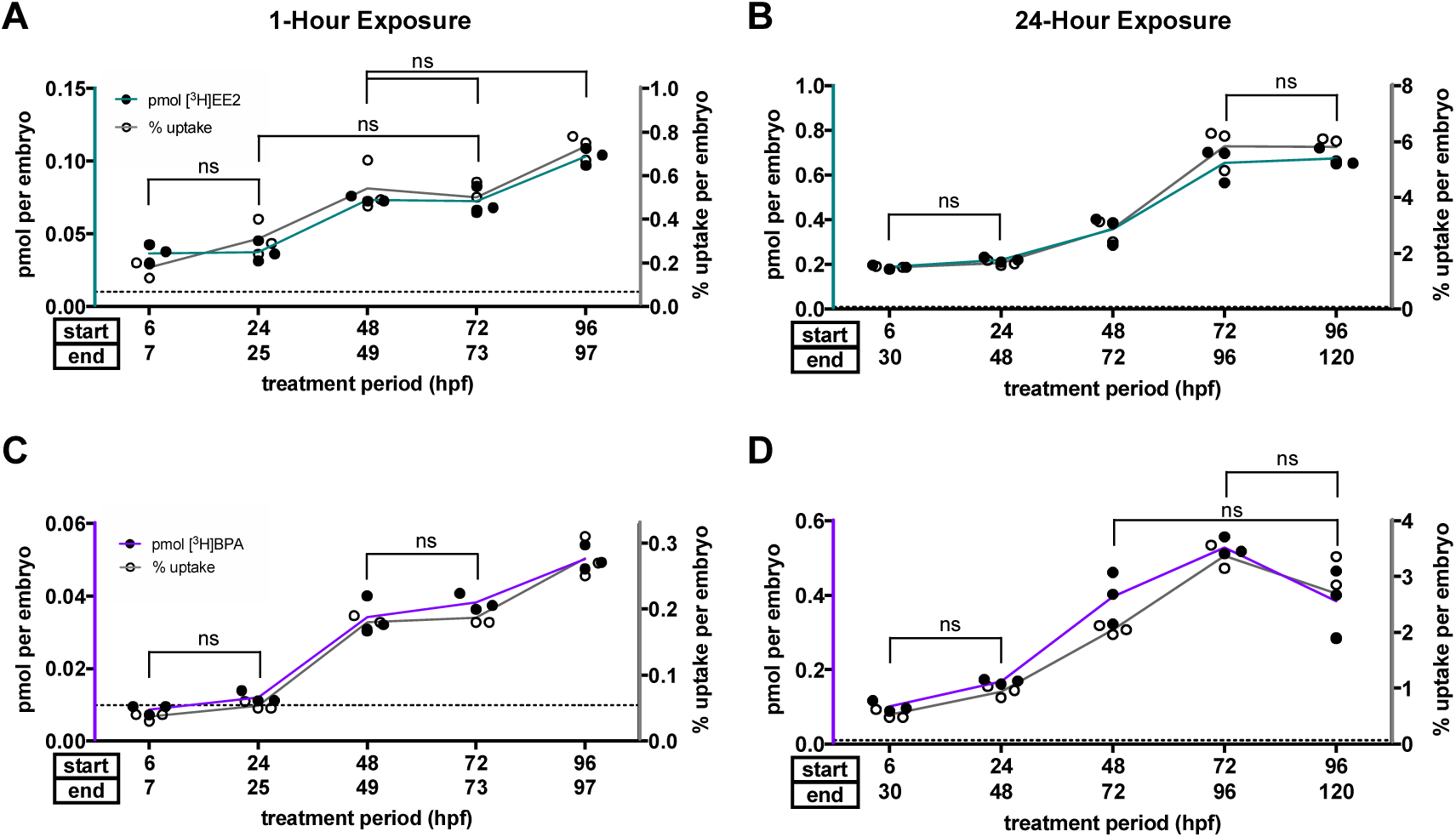
EE2 and BPA uptake depend on developmental stage in zebrafish. **(A, B)** Embryos and larvae were exposed to 5 nM [^3^H]EE2 **(A,B)** or 10 nM [^3^H]BPA **(C,D)** for one hour **(A,C)** or 24 hours **(B,D)** starting at five different developmental stages between 6 and 96 hours post fertilization (hpf). Radioactivity was measured using a scintillation counter and used to calculate pmol per embryo (black circles) and percent uptake per embryo (white circles). Each circle represents the mean uptake from a single experiment (n=3-5) assaying 10 embryos per experiment. Horizontal dotted line represents the limit of detection (0.01 pmol). Lines connect the mean for each group. Brackets denote percent uptake comparisons that are *not* statistically significant (ns), all other comparisons are significant (one-way ANOVA, p < 0.05).

Mean [^3^H]BPA pmol uptake increased to a greater extent, with a 5.6-fold higher uptake when starting treatment at 96 hpf versus 6 hpf (Fig. 2C, Table S3; 6 hpf treatment = 0.009 ± 0.001 pmol (mean ± SD), 24 hpf = 0.012 ± 0.002 pmol, 48 hpf = 0.034 ± 0.005 pmol, 72 hpf = 0.038 ± 0.002 pmol, 96 hpf = 0.050 ± 0.003 pmol). As with [^3^H]EE2, there was no significant difference in uptake when comparing uptake of embryos when starting treatment at 6 vs. 24 hpf (one-way ANOVA, p = 0.6854) or 48 vs. 72 hpf (p = 0.5343), but all other comparisons were significantly increased when starting treatment in older versus younger embryos (Table S3).

Percent uptake was similarly increased at 1 hour for both compounds. For [^3^H]EE2, there was a 4.1-fold increase when starting treatment at 96 hpf versus 6 hpf (Fig. 2A, Table S3; 6 hpf treatment = 0.18 ± 0.040 pmol (mean ± SD), 24 hpf = 0.31 ± 0.082 pmol, 48 hpf = 0.54 ± 0.11 pmol, 72 hpf = 0.50 ± 0.070 pmol, 96 hpf = 0.73 ± 0.057 pmol). No significant change was seen when comparing treatment start at 6 vs. 24 hpf (one-way ANOVA, p = 0.2794), 24 vs. 72 hpf (p = 0.0742), 48 vs. 72 hpf (p = 0.9649), or 48 vs. 96 hpf (p = 0.0683), but all other comparisons were significantly increased in older versus younger embryos (Table S3). [^3^H]BPA percent uptake increased 7.5-fold when starting treatment at 96 hpf vs. 6 hpf (Fig. 2C, Table S3; 6 hpf treatment = 0.037 ± 0.006 pmol (mean ± SD), 24 hpf = 0.053 ± 0.006 pmol, 48 hpf = 0.18 ± 0.010 pmol, 72 hpf = 0.19 ± 0.012 pmol, 96 hpf = 0.277 ± 0.031 pmol). Uptake was significantly increased when treating older versus younger embryos for every developmental stage (Table S3) except when comparing 6 vs. 24 hpf (one-way ANOVA, p = 0.6975) and 48 vs. 72 hpf embryos (p = 0.9833).

When increasing exposure duration to 24 hours, [^3^H]EE2 mean pmol uptake of beginning exposure at 96 hpf was 3.6-fold higher than beginning at 6 hpf, consistent with 1 hour exposure. In contrast, pmol absorption plateaued following exposure beginning at 72 hpf or older (Fig. 2B, Table S4; 6 hpf treatment = 0.19 ± 0.008 pmol (mean ± SD), 24 hpf = 0.22 ± 0.011 pmol, 48 hpf = 0.36 ± 0.062 pmol, 72 hpf = 0.66 ± 0.077 pmol, 96 hpf = 0.68 ± 0.040 pmol). There was a significantly higher uptake in older versus younger embryos at each developmental stage (Table S4) except when comparing exposures beginning at 6 vs. 24 hpf (one-way ANOVA, p = 0.9118) and 72 vs. 96 hpf (p = 0.9857).

There was a relatively smaller increase in [^3^H]BPA pmol uptake with developmental stage compared to 1 hour exposure, with a 3.8-fold increase when exposure began at 96 hpf compared to 6 hpf. This increase plateaued when exposure began at 48 hpf and older (Fig. 2D, Table S4; 6 hpf treatment = 0.10 ± 0.014 pmol (mean ± SD), 24 hpf = 0.17 ± 0.006 pmol, 48 hpf = 0.40 ± 0.070 pmol, 72 hpf = 0.53 ± 0.025 pmol, 96 hpf = 0.38 ± 0.091 pmol). There was a significantly higher uptake when comparing exposure starting at 48 vs. 96 hpf (one-way ANOVA, p = 0.0448). For each other comparison, there was a significantly higher uptake when starting exposure in older versus younger embryos (Table S4) except when comparing 6 vs. 24 hpf (one-way ANOVA, p = 0.5497), 48 vs. 72 hpf (p = 0.0688), and 48 vs. 96 hpf (p = 0.9985), which had no significant change.

Percent uptake following 24 hour exposure was similarly increased with increasing developmental stage for both compounds. Mean percent [^3^H]EE2 uptake was increased by 3.9-fold when starting treatment in 96 hpf versus 6 hpf embryos with a plateau ≥ 72 hpf (Fig. 2B, Table S4; 6 hpf treatment = 1.49 ± 0.051% (mean ± SD), 24 hpf = 1.65 ± 0.096%, 48 hpf = 2.88 ± 0.4%, 72 hpf = 5.82 ± 0.75%, 96 hpf = 5.81 ± 0.44%). Comparisons between exposure start at older and younger developmental stages showed significantly higher uptake when starting exposure at older stages (Table S4) except when comparing 6 vs. 24 hpf (one-way ANOVA, p = 0.9897) and 72 vs. 96 hpf (p > 0.9999). [^3^H]BPA percent uptake increased 5.1-fold in 96 hpf versus 6 hpf embryos with a plateau in uptake ≥ 48-72 hpf (Fig. 2D, Table S4; 6 hpf treatment = 0.53 ± 0.081% (mean ± SD), 24 hpf = 0.94 ± 0.10%, 48 hpf = 2.04 ± 0.080%, 72 hpf = 3.38 ± 0.21%, 96 hpf = 2.70 ± 0.75%). Uptake was significantly increased when starting treatment in older versus younger embryos (Table S4) except when comparing 6 vs. 24 hpf (one-way ANOVA, p = 0.6223), 48 vs. 96 hpf (0.2288), and 72 vs. 96 hpf (p = 0.2076).

### EE2 and E2 uptake exceed BPA uptake at multiple developmental stages

We next compared [^3^H]EE2 and [^3^H]BPA uptake at each developmental stage to [^3^H]E2 uptake previously reported(Souder and Gorelick, 2017). Percent uptake was used for all comparisons, as [^3^H]BPA required a higher exposure dose (10 nM) than [^3^H]EE2 and [^3^H]E2 (5 nM) for absorption above the limit of detection. Statistical significance was determined by comparing the mean log-value of the fold-change for three independent experiments to zero.

When comparing percent [^3^H]EE2 uptake following 1 hour exposure, there was a modest increase relative to [^3^H]E2 at embryos 6-48 hpf (Fig. 3C, Table S5; fold-change at 6 hpf = 1.72 ± 0.39 (mean ± SD), 24 hpf = 1.27 ± 0.33, 48 hpf = 1.41 ± 0.30, 72 hpf = 0.84 ± 0.12, 96 hpf = 0.81 ± 0.067), but no developmental stage had a statistically significant change (Table S5). When comparing [^3^H]BPA uptake to [^3^H]E2 uptake, there was a statistically significant decrease in uptake at each developmental stage (Fig. 3C, Table S5; one-way ANOVA; fold-change at 6 hpf = 0.36 ± 0.056, p = 0.0001; 24 hpf = 0.22 ± 0.023, p = 0.0001; 48 hpf = 0.47 ± 0.030, p = 0.0005; 72 hpf = 0.31 ± 0.023, p = 0.0001; 96 hpf = 0.31 ± 0.031, p = 0.0001).

**Figure 3.**
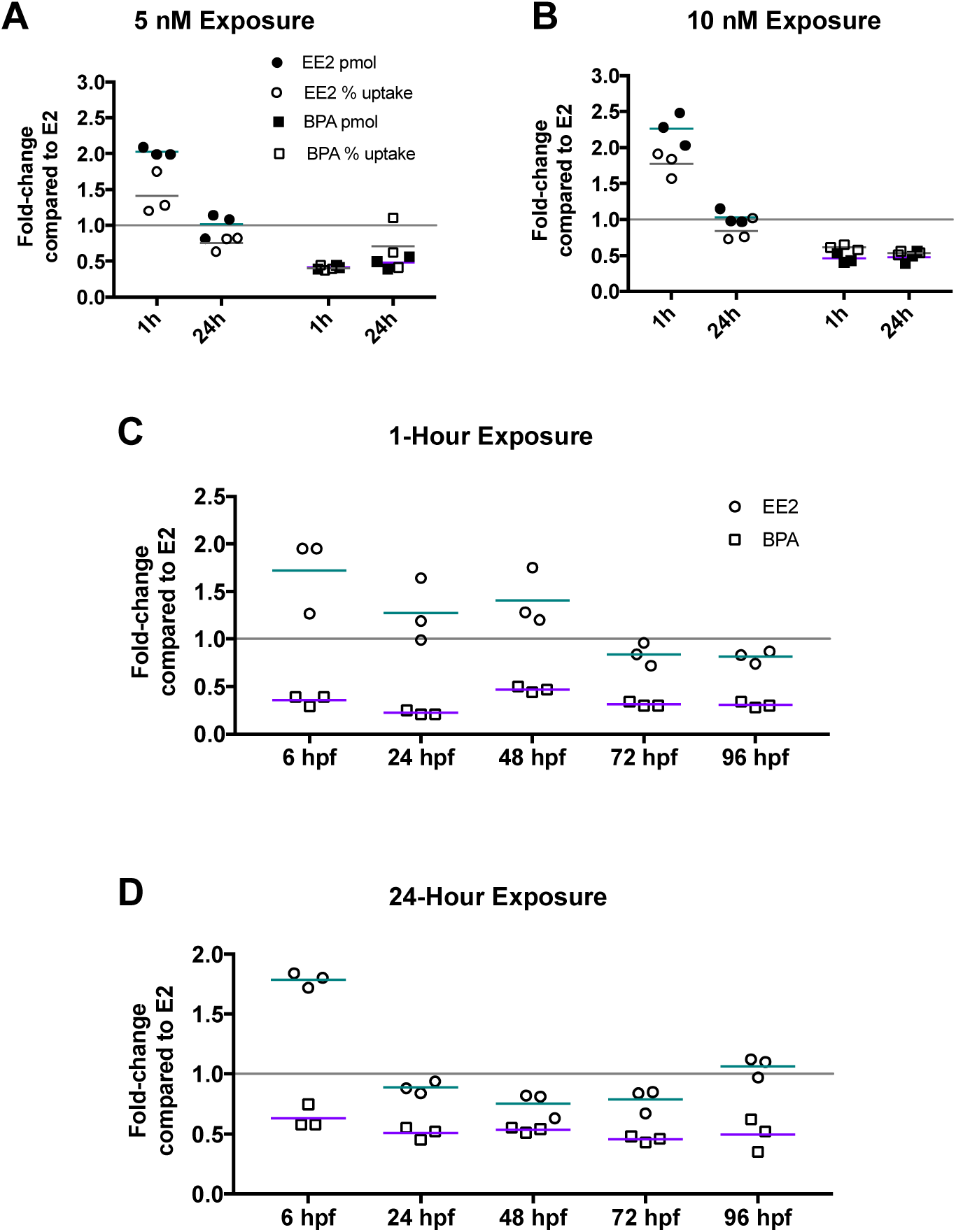
BPA uptake is less than E2 and EE2 uptake. **(A,B)** EE2 and BPA uptake compared to E2 uptake in 48 hours post fertilization (hpf) embryos exposed to 5 nM **(A)** or 10 nM **(B)** of each compound for 1 or 24 hours. **(C,D)** EE2 and BPA percent uptake compared to E2 in 6-96 hpf zebrafish exposed to 5 nM [^3^H]EE2 or [^3^H]E2 or 10 nM [^3^H]BPA for one hour **(C)** or 24 hours **(D)**. Fold-change compared to E2 was calculated by dividing the average pmol or percent uptake of [^3^H]EE2 or [^3^H]BPA (n=3 experiments, 10 embryos per experiment) by the pmol or percent uptake of [^3^H]E2 at each exposure condition. Horizontal lines represent the mean fold change in pmol (teal/purple) or percent uptake (gray). Horizontal gray line at y=1 represents no change in uptake compared to E2. [^3^H]E2 data from Souder and Gorelick, 2017.

With 24 hour exposure, we observed a statistically significant increase in [^3^H]EE2 percent uptake relative to [^3^H]E2 uptake in the 6 hpf group (one-way ANOVA, p = 0.0022), while older embryos demonstrated percent uptake similar to [^3^H]E2 (Fig. 3D, Table S5; fold-change at 6 hpf = 1.79 ± 0.062, 24 hpf = 0.89 ± 0.050, 48 hpf = 0.75 ± 0.11, 72 hpf = 0.79 ± 0.10, 96 hpf = 1.06 ± 0.081). Results with 24 hour exposure with [^3^H]BPA were consistent with the 1 hour exposure time, with an approximately 2-fold decrease in uptake at each developmental stage, though the decrease at 6 hpf was not statistically significant (Fig. 3D, Table S5; one-way ANOVA; fold-change at 6 hpf = 0.63 ± 0.097, p = 0.099; 24 hpf = 0.51 ± 0.051, p = 0.014; 48 hpf = 0.53 ± 0.021, p = 0.023; 72 hpf = 0.46 ± 0.025, p = 0.0059; 96 hpf = 0.50 ± 0.14, p = 0.0097). Therefore, we conclude that [^3^H]EE2 uptake was similar to [^3^H]E2 uptake, while [^3^H]BPA uptake was reduced compared to [^3^H]E2 uptake.

### Inhibition of zebrafish P-gp orthologue Abcb4 does not affect EED uptake

To test the idea that EEDs and their metabolites are transported by ABC transporters *in vivo,* we used our uptake assay as a proxy to measure efflux of [^3^H]E2 and [^3^H]BPA. We first asked whether treating embryos with a non-specific ABC transporter inhibitor known to inhibit Abcb4 (Fischer et al., 2013), CsA, would increase radioactivity of treated embryos following exposure via decreased efflux. Prior to experiments with [^3^H]E2, we confirmed that CsA could block efflux of a fluorescent substrate of Abcb4, rhodamine B (RhB), in 96 hpf embryos following 24 hour treatment. We found an increase in fluorescence with CsA and RhB co-treated embryos versus RhB-only treated embryos, confirming the efficacy of CsA at this developmental stage (Fig. 3B, one-way ANOVA; RhB = 1504 ± 40.44 units, RhB + CsA = 2445 ± 65.8 units, p = 0.0002) and consistent with previously published results in zebrafish (Fischer et al., 2013). We next tested whether the presence of E2 affected CsA activity by co-treating embryos with CsA, RhB, and non-radioactive E2. We did not observe a change in fluorescence when adding E2 at the same exposure concentration used for [^3^H] uptake studies (Fig. 4A, one-way ANOVA; RhB + E2 = 1586 ± 92.05 units, p = 0.7814 vs. RhB; RhB + CsA + E2 = 2488 ± 123.5, p = 0.9571 vs. RhB + CsA).

**Figure 4.**
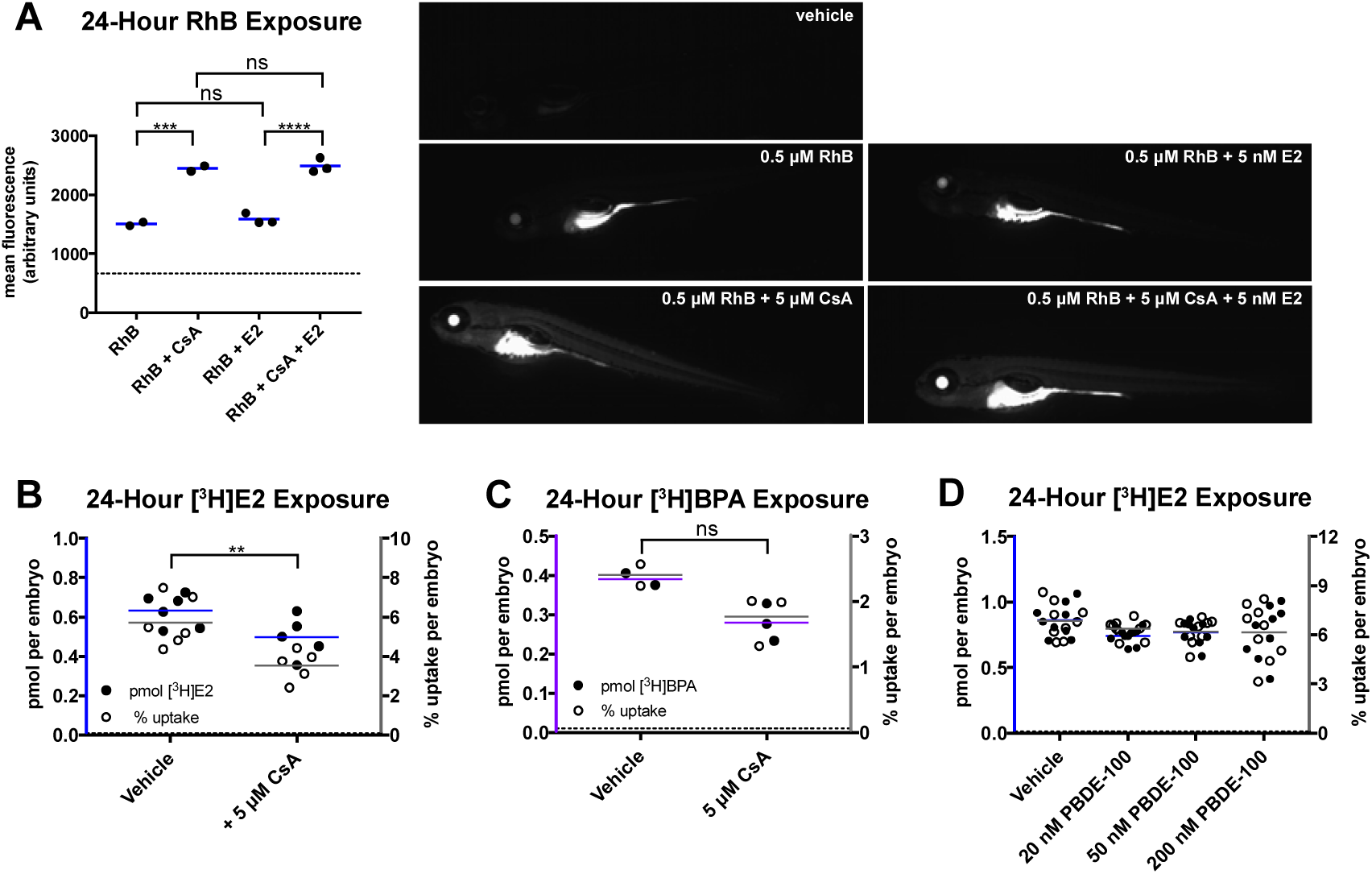
E2 uptake is not affected by ABC transporter inhibition. **(A)** Embryos were exposed at 96 hpf to vehicle or 0.5 μM rhodamine B (RhB) with or without 5 μM cyclosporin A (CsA) and non-radioactive 5 nM E2 for 24 hours in the dark. Embryos were imaged following treatment and mean integrated density of fluorescence for whole embryos was measured (mean fluorescence). Each circle on the graph represents the mean fluorescence from a single experiment (n=2-3 experiments, 10 embryos per experiment), horizontal blue line represents the mean. Horizontal dotted line at y=666 represents mean fluorescence of vehicle-treated embryos (n=10 embryos). Representative fluorescence images for each group are shown to the right of the graph. CsA treatment reduced RhB efflux, indicated by significantly increased RhB fluorescence. E2 exposure had no signficant affect on CsA-dependent RhB efflux (oneway ANOVA, ns=not significant, ^∗∗∗^p<0.001, ^∗∗∗∗^p<0.0001). Fluorescence of each group was significantly increased compared to vehicle (p<0.05). **(B)** Embryos at 96 hpf were exposed to 5 nM [^3^H]E2 with or without 5 μM CsA for 24 hours. Radioactivity was measured using a scintillation counter and used to calculate pmol per embryo (black circles) and percent uptake per embryo (white circles). Horizontal blue lines represent mean pmol uptake, gray lines represent mean percent uptake, n= 5-6 experiments, 10 embryos per experiment. CsA failed to increase uptake of [^3^H]E2. Instead, percent and pmol uptake were slightly decreased with the addition of CsA (unpaired Student’s t-test, ^∗∗^p<0.01 percent uptake, p<0.05 pmol uptake, CsA + E2 vs vehicle + E2). **(C)** Embryos at 96 hpf were exposed to 10 nM [^3^H]BPA with or without 5 μM CsA for 24 hours. Radioactivity was measured using a scintillation counter and used to calculate pmol per embryo (black circles) and percent uptake per embryo (white circles). Horizontal purple lines represent mean pmol uptake, gray lines represent mean percent uptake, n=2-3 experiments, 10 embryos per experiment. Percent [^3^H]BPA uptake was not significantly changed with the addition of CsA (unpaired Student’s t-test, ns not significant, p≥0.05). **(D)** Embryos at 72 hpf were exposed to 5 nM [^3^H]E2 together with vehicle or with PBDE-100 (20 - 200 nM). Radioactivity was measured using a scintillation counter and used to calculate pmol per embryo (black circles) and percent uptake per embryo (white circles). Each circle represents a single embryo, n=10 embryos per treatment. Horizontal blue lines represent mean pmol uptake, gray lines represent mean percent uptake. No significant change was found between groups (one-way ANOVA, p≥0.05). B-D, Horizontal dotted line represents the limit of detection (0.01 pmol).

To then test whether CsA inhibited [^3^H]E2 efflux, resulting in increased embryo radioactivity, we co-exposed embryos to CsA and [^3^H]E2. With 24 hour exposure at 96 hpf, pmol uptake was significantly decreased in CsA-treated embryos (Fig. 4B; Unpaired Student’s t-test; vehicle = 0.6336 ± 0.08162 pmol (mean ± SD), CsA = 0.4978 ± 0.1034 pmol, p = 0.0374). Similarly, percent uptake was significantly decreased in CsA-treated embryos (Fig. 4B; Unpaired Student’s t-test; vehicle = 5.72 ± 1.251% (mean ± SD), CsA = 3.534 ± 0.7851%, p = 0.0082). This suggests that CsA does not affect E2 transport, or that CsA modestly inhibits E2 uptake, resulting in reduced radioactivity.

We also tested the ability of CsA to block the efflux of BPA, a non-steroidal EED. Following 24 hour [^3^H]BPA exposure at 96 hpf, pmol BPA uptake was not significantly different in CsA-treated embryos compared to vehicle-treated embryos (Fig. 4C; Unpaired Student’s t-test; vehicle = 0.3909 ± 0.02145 pmol (mean ± SD), CsA = 0.2797 ± 0.04725 pmol, p = 0.06). Percent uptake was also not significantly different in CsA-treated embryos (Fig. 4C; Unpaired Student’s t-test; vehicle = 2.405 ± 0.2333 % (mean ± SD), CsA = 1.773 ± 0.3927 %, p = 0.14). This suggests that CsA does not affect BPA transport.

Concurrently, we treated zebrafish embryos with [^3^H]E2 and the POP 2,2’,4,4’,6- pentabromodiphenyl ether (PBDE-100), with the hypothesis that PBDE-100 would inhibit [^3^H]E2-conjugate efflux via P-gp or other ABC transporters, resulting in increased embryo radioactivity following exposure to E2 and PDBE-100 compared to E2 alone. Consistent with our CsA results, we found that there was no significant change in pmol uptake compared to vehicle at three concentrations of PBDE-100 (20 nM, 50 nM, 200 nM) (Fig. 4D; vehicle = 0.8604 ± 0.124 pmol (mean ± SD), 20 nM PBDE-100 = 0.7393 ± 0.05661 pmol, 50 nM PBDE-100 = 0.7677 ± 0.08959 pmol, 200 nM PBDE-100 = 0.7706 ± 0.1999 pmol). This suggests that co-exposure to PBDE-100 does not affect E2 transport.

## DISCUSSION

### Comparing uptake of estrogen-like EEDs

When comparing the uptake of different estrogen-like EEDs, we expected structurally similar EEDs like E2 and EE2 to have similar uptake. Since BPA is a non-steroidal estrogen and structurally distinct from E2, we expected relatively lower uptake with this compound. Our results were consistent with these hypotheses, as EE2 and E2 uptake were generally similar, while BPA uptake was lower than either EE2 or E2 uptake.

EE2 is a known environmental contaminant that may have adverse effects on aquatic wildlife, particularly during development (Bhandari et al., 2015; Santos et al., 2014; Volkova et al., 2015). Its structural similarity to E2, the primary circulating estrogen in humans, in conjunction with its high affinity for estrogen receptor alpha (ERα)(Pinto et al., 2014), makes EE2 relevant for toxicity assays in aquatic animals as well as animal models of human toxicity and teratogenicity. As a steroidal estrogen, EE2 is assumed to diffuse through cell membranes and be absorbed by lipophilic tissues, such as the embryonic yolk. We previously reported that, in embryos younger than 72 hpf, approximately 40-60% of exogenous E2 is absorbed by the yolk (Souder and Gorelick, 2017). We also found that the presence of the chorion did not significantly affect E2 uptake (Souder and Gorelick, 2017). We expect these results to hold true for EE2. For example, if 24 hpf embryos are exposed to E2 for 24 hours, we estimate that half the E2 that is taken up by the embryo will be deposited in the yolk. Extrapolating these results to EE2, which exhibited 1.6% uptake into the entire embryo, we predict that 0.8% of exogenous EE2 will be deposited in the yolk. This suggests that higher concentrations of EE2 are required earlier in development when the yolk comprises a larger proportion of the embryo. Acute treatments at these stages may also have longer term effects since EE2 absorbed into the yolk is available to diffuse into the embryo as it develops. The similar uptake results obtained with E2 and EE2 support the hypothesis that steroidal estrogens are absorbed equivalently by zebrafish embryos. Consequently, we speculate that the E2 and EE2 uptake properties are broadly applicable to any steroidal EED. Uptake of specific compounds can be confirmed using our isotopic assay. Finally, the similarity in E2 uptake, as an endogenous estrogen, and EE2 uptake, as a synthetic estrogen, suggests that endogenous estradiol levels do not limit E2 uptake at the levels present in zebrafish embryos and larvae. This suggests there are low levels of E2 present in the embryo, insufficient to affect passive diffusion gradients, or that there is some transport mechanism that is able to augment E2 uptake in embryos.

Comparing BPA uptake to E2 uptake can further inform future studies using nonsteroidal EEDs. While research testing the effects of BPA on development are abundant, many of these studies use environmentally-relevant low nanomolar concentrations of BPA in treatment water (Kinch et al., 2015; Kinch et al., 2016; Saili et al., 2012; Wu et al., 2017). Based on our results, the amount of BPA available to the embryo is substantially less than the reported EC_50_ of BPA for the zebrafish estrogen receptors ERα, 599 nM, ERβ1, 18.9 μM, and ERβ2, 3.8 μM (Pinto et al., 2014). For example, in 2015, Kinch and colleagues reported altered neurogenesis in embryos exposed to 6.8 nM BPA from 0-5 dpf. Based on our analysis of 10 nM exposure from 96-120 hpf, this corresponds to a concentration of 0.18 nM (2.7% uptake), well below the above listed EC_50_ values. Kinch and colleagues further reported that E2 treatment did not reproduce the effects of BPA, and that an androgen receptor antagonist, flutamide, was able to block the effects of BPA. This suggests that BPA may act via receptors other than estrogen receptors to elicit observed phenotypes. Another explanation for BPA effects at nanomolar concentrations is that BPA prevents the binding of endogenous estrogens to their receptors. As the EC_50_ of endogenous estrogens like E2 are hundreds-to thousands-fold greater than BPA (Pinto et al., 2014), however, it is unlikely that BPA displaces these estrogens at exposure concentrations below its EC_50_. Therefore, effects on aquatic model systems, like zebrafish, following exposure to low nanomolar concentrations of BPA may be due to non-estrogenic effects of BPA.

Overall, these results provide important reference information for exposing zebrafish embryos and larvae to environmentally relevant estrogen-like chemicals. Since percent uptake remains constant with increasing exposure concentration for the three EEDs described here, the percent uptake for the exposure conditions we tested can be used to estimate the concentration of a compound required in the water to achieve a specific concentration *in vivo.* Additionally, less severe phenotypes observed following exposure to BPA versus E2 may be due to decreased chemical availability rather than decreased efficacy. Therefore, comparisons of the effects of such compounds may require determination of *in vivo* levels.

### Comparison to uptake studies in other fish species

One study used [^3^H]EE2 and [^3^H]BPA to quantify uptake into fertilized medaka (*Oryzias latipes*) eggs (Bhandari et al., 2015). For [^3^H]EE2, they observed an uptake of 4.05 fmol/mg/egg following 24 hour 0.17 nM exposure. Assuming each egg is approximately 1 mg, this value is nearly 50-fold lower than the uptake we observed in 6 hpf embryos following 24 hour 5 nM exposure, though this discrepancy could be attributed to the 30fold lower concentration used in their study. They observed [^3^H]BPA uptake of 0.125 pmol/mg/egg following 24 hour 44 nM expousre, which is comparable to the 0.1005 pmol we observed following 24 hour 10 nM exposure.

### Comparison to uptake studies using chromatography and mass spectrometry

In one study using high-performance liquid chromatography coupled to mass spectrometry (HPLC-MS) to measure BPA uptake into zebrafish embryos, there was a 0.02 μg/kg uptake observed following 1 nM exposure from 8-58 hpf (Saili et al., 2012). Assuming a mass of 0.291 mg per embryo (Kantae et al., 2016), this corresponds to approximately 0.025 fmol BPA per embryo. The most relevant comparison to the Saili exposure conditions that we tested is a 10 nM exposure from 24-48 hpf. We observed an uptake of 0.168 pmol, which could be estimated as 0.0168 pmol with 1 nM exposure (an exposure concentration that was below our limit of detection). Two differences between our study and the Saili study are the standard curve and the pooling of embryos. First, Saili and colleagues extrapolated uptake with 1 nM exposure from a standard curve produced using much higher exposure concentrations of BPA (1-100 μM). Our assay provides a means of measurement that is more sensitive, with detectable levels of BPA following exposure to 5 nM concentration, rather than 1 μM. Second, Saili and colleagues used pools of 50 embryos to quantify uptake at each exposure concentration. Our assay is able to measure single embryos, which is more useful for screening compounds that affect BPA uptake.

Wu and colleagues also used HPLC-MS to measure BPA uptake in zebrafish embryos from 4-72 hpf with multiple exposure concentrations (Wu et al., 2017). Comparing their 1 μg/L exposure (4.38 nM) to our 5 nM treatment from 48-72 hpf yields similar results, with 0.196 pmol/embryo via HPLC, and 0.190 pmol/embryo via our isotopic assay. This provides validity for the accuracy of our assay, despite its simplicity compared to the sophisticated instruments required for HPLC-MS analysis.

### Effect of ABC transporter inhibition on EED uptake and toxicity

The isotopic uptake assay can test the effect of environmental chemicals on EED uptake. Persistent organic pollutants (POPs) can inhibit ABC transporters, which may contribute to POP toxicity in aquatic animals by preventing efflux of small molecules, such as EEDs, that are normally effluxed by ABC transporters (Nicklisch et al., 2016). Estradiol is assumed to freely diffuse through cell membranes, but its less lipophilic conjugates, such as 17-β-*d*-E2-glucuronide, are transported by ABC transporters like Pgp (Huang et al., 1998). The zebrafish paralogue of P-gp, encoded by the *abcb4* gene, transports known substrates of human P-gp, including cyclosporin A (CsA), vinblastine, and rhodamine B (RhB)(Fischer et al., 2013). Abcb4 is expressed ubiquitously early in zebrafish development, and becomes localized to the gut by 120 hpf (Fischer et al., 2013). Further, the enzymes that metabolize E2—such as UDP-glucuronosyltransferases—are expressed early in development in zebrafish (Christen and Fent, 2014), therefore E2 is likely metabolized in zebrafish embryos and could be effluxed by Abcb4. We found, however, that co-treatment with the Pg-p inhibitor CsA modestly decreased [^3^H]E2 uptake, and that PBDE-100 had no effect on uptake. One possible explanation for the decrease in uptake observed with CsA treatment is that the transporters blocked by CsA are normally involved in augmenting [^3^H]E2 uptake, rather than controlling its efflux. Additionally, the decrease in percent uptake with CsA cotreatment is relatively low (2.2%), and may not be biologically relevant. When adjusting water concentration to account for the small percentage absorbed by the embryos, there is an effective concentration of 286 pM without CsA, and 176 pM with CsA treatment. Both of these values exceed the EC_50_ of E2 for ERα (77 pM), ERβ1 (39 pM), and ERβ2 (118 pM)(Pinto et al., 2014).

We performed the same experiment using [^3^H]BPA, to test the effect of CsA-mediated ABC transporter inhibition on a non-steroidal EED. It has been suggested by *in vitro* assays that BPA transport by ABC transporters occurs in a species-specific manner (Mazur et al., 2012). It is not well-understood how BPA is normally transported in developing zebrafish embryos. Our results suggest that BPA is not effluxed by CsA-responsive transporters in zebrafish embryos 96-120 hpf.

It is possible that the concentrations of E2 and BPA we used in our experiments are below the concentrations required to stimulate Abcb4 activity. Future studies could treat embryos with a mixture of non-radioactive and radioactive compounds to increase exposure concentrations, and determine if higher concentrations are able to inhibit Abcb4 or other ABC transporters. Further, the promiscuity of CsA for ABC transporters may cause inhibition of transporters with opposing directionality in cellular membranes, facilitating efflux versus influx. To identify the role of specific transporters involved in E2 or BPA transport, embryos could be treated with isoform-specific antagonists of the ABC transporters. This could also be accomplished with the generation of mutant zebrafish lines with non-functional transporters.

### Conclusions and future directions

The results presented here confirm the utility of the tritium-based isotopic uptake assay for sensitive measurement of structurally-diverse EEDs *in vivo.* In contrast to HPLC and mass spectrometry approaches, the isotopic uptake assay requires only a scintillation counter, an instrument to which many labs have access and which requires little specialized training to operate. Additionally, the isotopic assay does not require any chemical extraction procedures. We also present this assay as a screening tool for identification of chemicals that affect EED transport. Future studies could expand upon these results to test the uptake of non-estrogenic EEDs and other steroid hormones, like androgens and progesterone, and identify proteins involved in compound transport via pharmacologic or genetic manipulation. Our results provide important information regarding the comparison of toxicity between structurally-distinct EEDs and will inform future studies using these compounds to investigate mechanisms of EED toxicity.

## SUPPLEMENTARY DATA

Supplementary figure S1 and tables S1–S6 supplied below Figures.

## ACKNOWLEDGMENTS

We thank Stephen G. Aller for sharing chemicals and insights regarding P-gp function, Charles N. Falany for use of materials and advice on isotopic uptake, and Susan Farmer and the UAB zebrafish facility staff for zebrafish husbandry and maintenance.

### FUNDING INFORMATION

This work was supported by National Institute of Environmental Health Sciences [R01ES026337] and National Institute of General Medical Sciences [T32GM008361].

## SUPPLEMENTARY DATA

**Figure S1.**
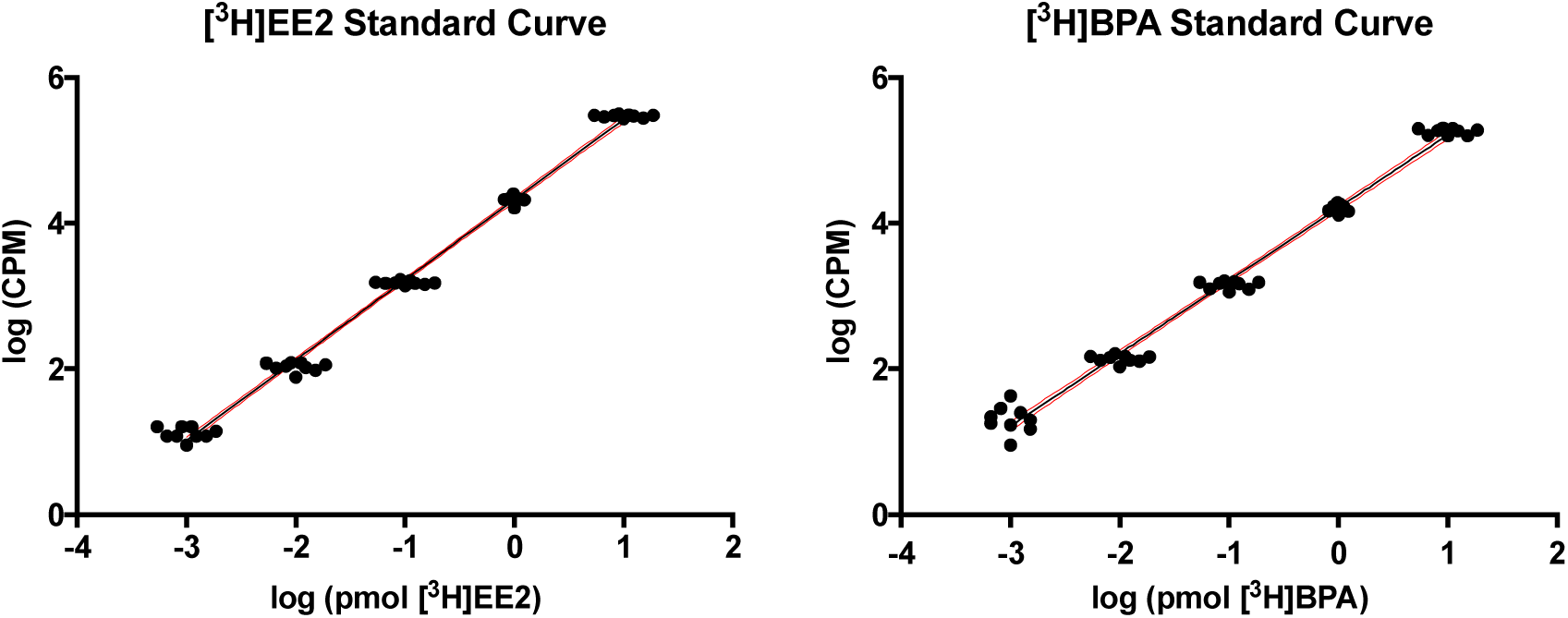
Standard curve for uptake calculations. **(A-B)** The radioactivity of 10 μL of a known concentration of [^3^H]EE2 **(A)** or [^3^H]BPA **(B)** was measured on a scintillation counter in triplicate on three different days for a total of 9 measurements per dose (black circles). A linear relationship was found when performing a power analysis of the mean of each concentration. **(A)** Y = 1.101^∗^X + 4.326 where X = log(pmol [^3^H]EE2) and Y = log(CPM). r^2^ = 0.9972 (black line; unweighted, best-fit linear regression line). **(B)** Y = 0.998^∗^X + 4.212 where X = log(pmol [^3^H]BPA) and Y = log(CPM). r^2^ = 0.9939 (black line; unweighted, best-fit linear regression line). Outer red lines represent 95% CI.

**Table S1.** Fold-change vs. E2 pmol and percent uptake following 1-hour exposure. Fold-change was calculated by dividing pmol [^3^H]EE2 or [^3^H]BPA by published pmol absorption [^3^H]E2 (Souder and Gorelick, 2017). Adjusted p-values were determined by one-way ANOVA with Dunnett’s test for multiple comparisons to compare log values of the fold-change to zero. ^∗^p<0.05, ^∗∗^p<0.01, ns not significant (p≥0.05).

| 1-Hour Exposure |  |  |  |  |  |
| --- | --- | --- | --- | --- | --- |
| pmol [<sup>3</sup>H]EE2 |  |  |  |  |  |
| Exposure Concentration (nM) | mean [<sup>3</sup>H]E2 (pmol) | mean [<sup>3</sup>H]EE2 (pmol) | log(fold-change) | adjusted p-value | summary |
| 5 | 0.0364 | 0.0736 | 0.3060 | 0.0060 | ** |
| 10 | 0.0746 | 0.1689 | 0.3533 | 0.0028 | ** |
| % uptake [<sup>3</sup>H]EE2 |  |  |  |  |  |
| Exposure Concentration (nM) | mean [<sup>3</sup>H]E2 (%) | mean [<sup>3</sup>H]EE2 (%) | log(fold-change) | adjusted p-value | summary |
| 5 | 0.3833 | 0.5400 | 0.1431 | 0.7113 | ns |
| 10 | 0.2933 | 0.5200 | 0.2473 | 0.0071 | ** |
| pmol [<sup>3</sup>H]BPA |  |  |  |  |  |
| Exposure Concentration (nM) | mean [<sup>3</sup>H]E2 (pmol) | mean [<sup>3</sup>H]BPA (pmol) | log(fold-change) | adjusted p-value | summary |
| 5 | 0.0364 | 0.0149 | -0.3842 | 0.0015 | ** |
| 10 | 0.0746 | 0.0341 | -0.3405 | 0.0035 | ** |
| % uptake [<sup>3</sup>H]BPA |  |  |  |  |  |
| Exposure Concentration (nM) | mean [<sup>3</sup>H]E2 (%) | mean [<sup>3</sup>H]BPA (%) | log(fold-change) | adjusted p-value | summary |
| 5 | 0.3833 | 0.1533 | -0.3991 | 0.0662 | ns |
| 10 | 0.2933 | 0.1800 | -0.2128 | 0.0163 | * |

**Table S2.** Fold-change vs. E2 pmol and percent uptake following 24-hour exposure. Fold-change was calculated by dividing pmol [^3^H]EE2 or [^3^H]BPA by published pmol absorption [^3^H]E2 (Souder and Gorelick, 2017). Adjusted p-values were determined by one-way ANOVA with Dunnett’s test for multiple comparisons to compare log values of the fold-change to zero. ^∗^p<0.05, ^∗∗^p<0.01, ns not significant (p≥0.05).

| 24-Hour Exposure |  |  |  |  |  |
| --- | --- | --- | --- | --- | --- |
| pmol [<sup>3</sup>H]EE2 |  |  |  |  |  |
| Exposure Concentration (nM) | mean [<sup>3</sup>H]E2 (pmol) | mean [<sup>3</sup>H]EE2 (pmol) | log(fold-change) | adjusted p-value | summary |
| 5 | 0.3544 | 0.3582 | -0.0004 | 0.9999 | ns |
| 10 | 0.8263 | 0.8538 | 0.0129 | 0.9989 | ns |
| % uptake [<sup>3</sup>H]EE2 |  |  |  |  |  |
| Exposure Concentration (nM) | mean [<sup>3</sup>H]E2 (%) | mean [<sup>3</sup>H]EE2 (%) | log(fold-change) | adjusted p-value | summary |
| 5 | 3.8200 | 2.8800 | -0.1261 | 0.7844 | ns |
| 10 | 3.8067 | 3.1970 | -0.0824 | 0.4378 | ns |
| pmol [<sup>3</sup>H]BPA |  |  |  |  |  |
| Exposure Concentration (nM) | mean [<sup>3</sup>H]E2 (pmol) | mean [<sup>3</sup>H]BPA (pmol) | log(fold-change) | adjusted p-value | summary |
| 5 | 0.3544 | 0.1904 | -0.2723 | 0.0114 | * |
| 10 | 0.8263 | 0.3953 | -0.3235 | 0.0048 | ** |
| % uptake [<sup>3</sup>H]BPA |  |  |  |  |  |
| Exposure Concentration (nM) | mean [<sup>3</sup>H]E2 (%) | mean [<sup>3</sup>H]BPA (%) | log(fold-change) | adjusted p-value | summary |
| 5 | 3.8200 | 2.2717 | -0.1845 | 0.5311 | ns |
| 10 | 3.8067 | 2.0430 | -0.2706 | 0.0042 | ** |

**Table S3.** EE2 and BPA uptake with increasing developmental stage following 1-hour exposure. Mean 1 denotes the mean pmol or percent uptake of the developmental stage compared to (Start hpf). Mean 2 denotes the mean pmol or percent uptake of starting treatment at higher developmental stages (vs hpf). Adjusted p-values comparing the mean uptake at each developmental stage were determined via one-way ANOVA with Tukey’s test for multiple comparisons. ^∗^p<0.05, ^∗∗^p<0.01, ^∗∗∗^p<0.001, ^∗∗∗∗^p<0.0001, ns not significant (p≥0.05).

| 1-Hour Exposure |  |  |  |  |  |  |
| --- | --- | --- | --- | --- | --- | --- |
| pmol [3H]EE2 |  |  |  |  |  |  |
| Start (hpf) | mean 1 | vs (hpf) | mean 2 | mean 2 - mean 1 | adjusted p-value | summary |
| 6 | 0.0365 | 24 | 0.0376 | 0.0011 | 0.9996 | ns |
|  |  | 48 | 0.0736 | 0.0371 | 0.0003 | *** |
|  |  | 72 | 0.0723 | 0.0358 | 0.0004 | *** |
|  |  | 96 | 0.1033 | 0.0668 | <0.0001 | **** |
| 24 | 0.0376 | 48 | 0.0736 | 0.0360 | 0.0004 | *** |
|  |  | 72 | 0.0723 | 0.0347 | 0.0005 | *** |
|  |  | 96 | 0.1033 | 0.0657 | <0.0001 | **** |
| 48 | 0.0736 | 72 | 0.0723 | -0.0013 | 0.9989 | ns |
|  |  | 96 | 0.1033 | 0.0297 | 0.0018 | ** |
| 72 | 0.0723 | 96 | 0.1033 | 0.0310 | 0.0013 | ** |
| % uptake [3H]EE2 |  |  |  |  |  |  |
| Start (hpf) | mean 1 | vs (hpf) | mean 2 | mean 2 - mean 1 | adjusted p-value | summary |
| 6 | 0.1767 | 24 | 0.3100 | 0.1333 | 0.2794 | ns |
|  |  | 48 | 0.5400 | 0.3633 | 0.0012 | ** |
|  |  | 72 | 0.5000 | 0.3233 | 0.0030 | ** |
|  |  | 96 | 0.7333 | 0.5567 | <0.0001 | **** |
| 24 | 0.3100 | 48 | 0.5400 | 0.2300 | 0.0275 | * |
|  |  | 72 | 0.5000 | 0.1900 | 0.0742 | ns |
|  |  | 96 | 0.7333 | 0.4233 | 0.0004 | *** |
| 48 | 0.5400 | 72 | 0.5000 | -0.0400 | 0.9649 | ns |
|  |  | 96 | 0.7333 | 0.1933 | 0.0683 | ns |
| 72 | 0.5000 | 96 | 0.7333 | 0.2333 | 0.0254 | * |
| pmol [3H]BPA |  |  |  |  |  |  |
| Start (hpf) | mean 1 | vs (hpf) | mean 2 | mean 2 - mean 1 | adjusted p-value | summary |
| 6 | 0.0087 | 24 | 0.0121 | 0.0034 | 0.6854 | ns |
|  |  | 48 | 0.0341 | 0.0254 | <0.0001 | **** |
|  |  | 72 | 0.0381 | 0.0294 | <0.0001 | **** |
|  |  | 96 | 0.0502 | 0.0415 | <0.0001 | **** |
| 24 | 0.0121 | 48 | 0.0341 | 0.0220 | <0.0001 | **** |
|  |  | 72 | 0.0381 | 0.0260 | <0.0001 | **** |
|  |  | 96 | 0.0502 | 0.0381 | <0.0001 | **** |
| 48 | 0.0341 | 72 | 0.0381 | 0.0040 | 0.5343 | ns |
|  |  | 96 | 0.0502 | 0.0161 | 0.0006 | *** |
| 72 | 0.0381 | 96 | 0.0502 | 0.0121 | 0.0050 | ** |
| % uptake [3H]BPA |  |  |  |  |  |  |
| Start (hpf) | mean 1 | vs (hpf) | mean 2 | mean 2 - mean 1 | adjusted p-value | summary |
| 6 | 0.0367 | 24 | 0.0533 | 0.0167 | 0.6975 | ns |
|  |  | 48 | 0.1800 | 0.1433 | <0.0001 | **** |
|  |  | 72 | 0.1867 | 0.1500 | <0.0001 | **** |
|  |  | 96 | 0.2767 | 0.2400 | <0.0001 | **** |
| 24 | 0.0533 | 48 | 0.1800 | 0.1267 | <0.0001 | **** |
|  |  | 72 | 0.1867 | 0.1333 | <0.0001 | **** |
|  |  | 96 | 0.2767 | 0.2233 | <0.0001 | **** |
| 48 | 0.1800 | 72 | 0.1867 | 0.0067 | 0.9833 | ns |
|  |  | 96 | 0.2767 | 0.0967 | 0.0001 | *** |
| 72 | 0.1867 | 96 | 0.2767 | 0.0900 | 0.0003 | *** |

**Table S4.** EE2 and BPA uptake with increasing developmental stage following 24-hour exposure. Mean 1 denotes the mean pmol or percent uptake of the developmental stage compared to (Start hpf). Mean 2 denotes the mean pmol or percent uptake of starting treatment at higher developmental stages (vs hpf). Adjusted p-values comparing mean uptake at each developmental stage were determined via one-way ANOVA with Tukey’s test for multiple comparisons. ^∗^p<0.05, ^∗∗^p<0.01, ^∗∗∗^p<0.001, ^∗∗∗∗^p<0.0001, ns not significant (p≥0.05).

| 24-Hour Exposure |  |  |  |  |  |  |
| --- | --- | --- | --- | --- | --- | --- |
| pmol [3H]EE2 |  |  |  |  |  |  |
| Start (hpf) | mean 1 | vs (hpf) | mean 2 | mean 2 - mean 1 | adjusted p-value | summary |
| 6 | 0.1891 | 24 | 0.2222 | 0.0331 | 0.9118 | ns |
|  |  | 48 | 0.3582 | 0.1691 | 0.0107 | * |
|  |  | 72 | 0.6562 | 0.4671 | <0.0001 | **** |
|  |  | 96 | 0.6758 | 0.4867 | <0.0001 | **** |
| 24 | 0.2222 | 48 | 0.3582 | 0.1360 | 0.0388 | * |
|  |  | 72 | 0.6562 | 0.4340 | <0.0001 | **** |
|  |  | 96 | 0.6758 | 0.4536 | <0.0001 | **** |
| 48 | 0.3582 | 72 | 0.6562 | 0.2980 | 0.0001 | *** |
|  |  | 96 | 0.6758 | 0.3176 | <0.0001 | **** |
| 72 | 0.6562 | 96 | 0.6758 | 0.0196 | 0.9857 | ns |
| % uptake [3H]EE2 |  |  |  |  |  |  |
| Start (hpf) | mean 1 | vs (hpf) | mean 2 | mean 2 - mean 1 | adjusted p-value | summary |
| 6 | 1.4867 | 24 | 1.6467 | 0.1600 | 0.9897 | ns |
|  |  | 48 | 2.8800 | 1.3933 | 0.0175 | * |
|  |  | 72 | 5.8200 | 4.3333 | <0.0001 | **** |
|  |  | 96 | 5.8100 | 4.3233 | <0.0001 | **** |
| 24 | 1.6467 | 48 | 2.8800 | 1.2333 | 0.0354 | * |
|  |  | 72 | 5.8200 | 4.1733 | <0.0001 | **** |
|  |  | 96 | 5.8100 | 4.1633 | <0.0001 | **** |
| 48 | 2.8800 | 72 | 5.8200 | 2.9400 | <0.0001 | **** |
|  |  | 96 | 5.8100 | 2.9300 | <0.0001 | **** |
| 72 | 5.8200 | 96 | 5.8100 | -0.0100 | >0.9999 | ns |
| pmol [3H]BPA |  |  |  |  |  |  |
| Start (hpf) | mean 1 | vs (hpf) | mean 2 | mean 2 - mean 1 | adjusted p-value | summary |
| 6 | 0.1005 | 24 | 0.1680 | 0.0675 | 0.5497 | ns |
|  |  | 48 | 0.3953 | 0.2948 | 0.0003 | *** |
|  |  | 72 | 0.5285 | 0.4280 | <0.0001 | **** |
|  |  | 96 | 0.3833 | 0.2828 | 0.0005 | *** |
| 24 | 0.1680 | 48 | 0.3953 | 0.2273 | 0.0026 | ** |
|  |  | 72 | 0.5285 | 0.3605 | <0.0001 | **** |
|  |  | 96 | 0.3833 | 0.2153 | 0.0039 | ** |
| 48 | 0.3953 | 72 | 0.5285 | 0.1332 | 0.0688 | ns |
|  |  | 96 | 0.3833 | -0.0120 | 0.9985 | ns |
| 72 | 0.5285 | 96 | 0.3833 | -0.1452 | 0.0448 | * |
| % uptake [3H]BPA |  |  |  |  |  |  |
| Start (hpf) | mean 1 | vs (hpf) | mean 2 | mean 2 - mean 1 | adjusted p-value | summary |
| 6 | 0.5267 | 24 | 0.9400 | 0.4133 | 0.6223 | ns |
|  |  | 48 | 2.0433 | 1.5166 | 0.0026 | ** |
|  |  | 72 | 3.3767 | 2.8500 | <0.0001 | **** |
|  |  | 96 | 2.7000 | 2.1733 | 0.0001 | *** |
| 24 | 0.9400 | 48 | 2.0433 | 1.1033 | 0.0218 | * |
|  |  | 72 | 3.3767 | 2.4367 | <0.0001 | **** |
|  |  | 96 | 2.7000 | 1.7600 | 0.0008 | *** |
| 48 | 2.0433 | 72 | 3.3767 | 1.3334 | 0.0066 | ** |
|  |  | 96 | 2.7000 | 0.6567 | 0.2288 | ns |
| 72 | 3.3767 | 96 | 2.7000 | -0.6767 | 0.2076 | ns |

**Table S5.**
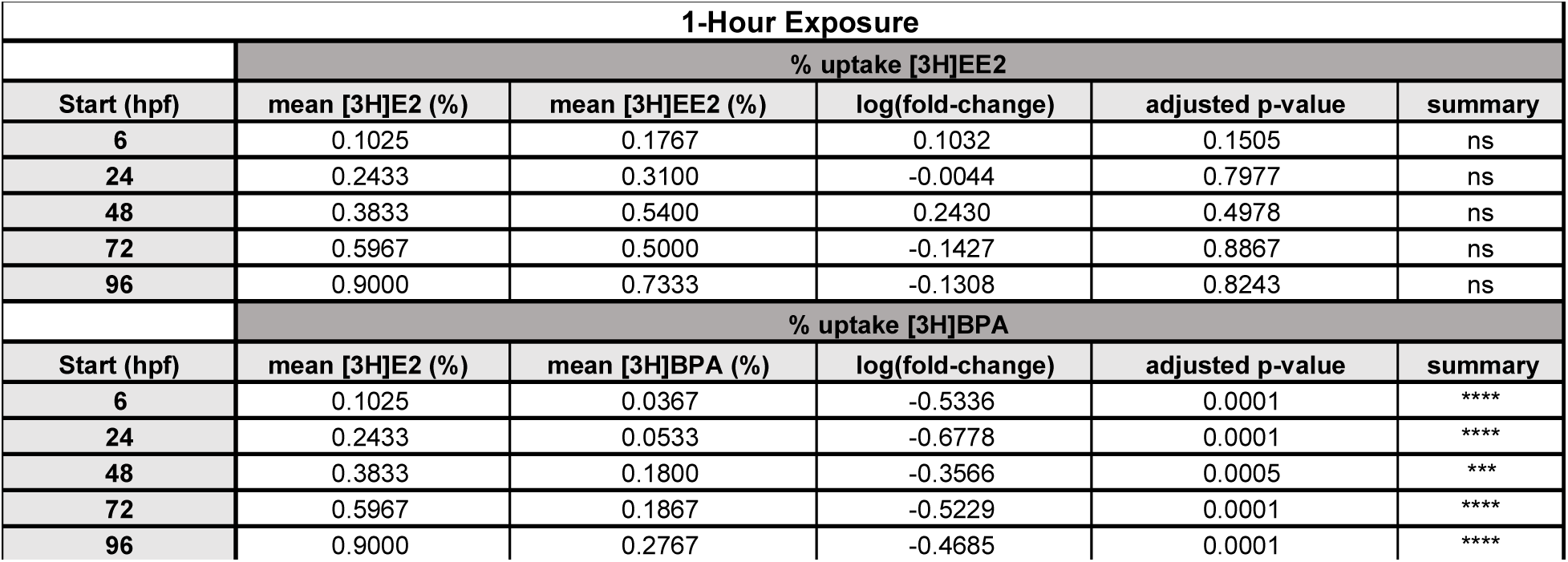
Fold-change vs E2 percent uptake with 1-hour exposure. Fold-change was calculated by dividing pmol [^3^H]EE2 or [^3^H]BPA by published pmol absorption [^3^H]E2.(Souder and Gorelick, 2017) Adjusted p-values were determined by one-way ANOVA with Dunnett’s test for multiple comparisons to compare log values of the fold-change to zero. ^∗∗∗^p<0.001, ^∗∗∗∗^p<0.0001, ns not significant (p≥0.05).

| 1-Hour Exposure |  |  |  |  |  |
| --- | --- | --- | --- | --- | --- |
| % uptake [<sup>3</sup>H]EE2 |  |  |  |  |  |
| Start (hpf) | mean [<sup>3</sup>H]E2 (%) | mean [<sup>3</sup>H]EE2 (%) | log(fold-change) | adjusted p-value | summary |
| 6 | 0.1025 | 0.1767 | 0.1032 | 0.1505 | ns |
| 24 | 0.2433 | 0.3100 | -0.0044 | 0.7977 | ns |
| 48 | 0.3833 | 0.5400 | 0.2430 | 0.4978 | ns |
| 72 | 0.5967 | 0.5000 | -0.1427 | 0.8867 | ns |
| 96 | 0.9000 | 0.7333 | -0.1308 | 0.8243 | ns |
| % uptake [<sup>3</sup>H]BPA |  |  |  |  |  |
| Start (hpf) | mean [<sup>3</sup>H]E2 (%) | mean [<sup>3</sup>H]BPA (%) | log(fold-change) | adjusted p-value | summary |
| 6 | 0.1025 | 0.0367 | -0.5336 | 0.0001 | **** |
| 24 | 0.2433 | 0.0533 | -0.6778 | 0.0001 | **** |
| 48 | 0.3833 | 0.1800 | -0.3566 | 0.0005 | *** |
| 72 | 0.5967 | 0.1867 | -0.5229 | 0.0001 | **** |
| 96 | 0.9000 | 0.2767 | -0.4685 | 0.0001 | **** |

**Table S6.** Fold-change vs E2 percent uptake with 24-hour exposure. Fold-change was calculated by dividing pmol [^3^H]EE2 or [^3^H]BPA by published pmol absorption [^3^H]E2 (Souder and Gorelick, 2017). Adjusted p-values were determined by one-way ANOVA with Dunnett’s test for multiple comparisons to compare log values of the fold-change to zero. ^∗^p<0.05, ^∗∗^p<0.01, ns not significant (p≥0.05).

| 24-Hour Exposure |  |  |  |  |  |
| --- | --- | --- | --- | --- | --- |
| % uptake [<sup>3</sup>H]EE2 |  |  |  |  |  |
| Start (hpf) | mean [<sup>3</sup>H]E2 (%) | mean [<sup>3</sup>H]EE2 (%) | log(fold-change) | adjusted p-value | summary |
| 6 | 0.8320 | 1.4867 | 0.2352 | 0.0022 | ** |
| 24 | 1.8625 | 1.6467 | -0.0757 | 0.7479 | ns |
| 48 | 3.8200 | 2.8800 | -0.0915 | 0.1101 | ns |
| 72 | 7.3800 | 5.8200 | -0.0706 | 0.1992 | ns |
| 96 | 5.4600 | 5.8100 | 0.0492 | 0.9778 | ns |
| % uptake [<sup>3</sup>H]BPA |  |  |  |  |  |
| Start (hpf) | mean [<sup>3</sup>H]E2 (%) | mean [<sup>3</sup>H]BPA (%) | log(fold-change) | adjusted p-value | summary |
| 6 | 0.8320 | 0.5267 | -0.2388 | 0.0988 | ns |
| 24 | 1.8625 | 0.9400 | -0.3468 | 0.0142 | * |
| 48 | 3.8200 | 2.0433 | -0.2924 | 0.0229 | * |
| 72 | 7.3800 | 3.3767 | -0.3188 | 0.0059 | ** |
| 96 | 5.4600 | 2.7000 | -0.2840 | 0.0097 | ** |

